# Unraveling membrane properties at the organelle-level with LipidDyn

**DOI:** 10.1101/2022.01.04.474788

**Authors:** Simone Scrima, Matteo Tiberti, Alessia Campo, Elisabeth Corcelle-Termeau, Delphine Judith, Mads Møller Foged, Knut Kristoffer Bundgaard Clemmensen, Sharon Tooze, Marja Jäättelä, Kenji Maeda, Matteo Lambrughi, Elena Papaleo

**Affiliations:** Cancer Structural Biology, Center for Autophagy, Recycling and Disease, Danish Cancer Society Research Center, 2100, Copenhagen, Denmark; Cancer Systems Biology, Section for Bioinformatics, Department of Health and Technology, Technical University of Denmark, Lyngby, Denmark; Cell Death and Metabolism, Center for Autophagy, Recycling and Disease, Danish Cancer Society Research Center, 2100, Copenhagen, Denmark; Institut Cochin, Inserm U1016-CNRS, UMR8104, Université de Paris, Paris, France; Molecular Cell Biology and Autophagy Laboratory, The Francis Crick Institute, London NW1 1AT, UK

**Author notes:** Corresponding authors: Elena Papaleo,; Matteo Lambrughi. These authors jointly supervised this work (Matteo Lambrughi, Elena Papaleo).

**Keywords:** molecular dynamics, lipid structure, lipidomics, organelles, protein-lipid interactions, autophagy

## Abstract

Cellular membranes are formed from many different lipids in various amounts and proportions depending on the subcellular localization. The lipid composition of membranes is sensitive to changes in the cellular environment, and their alterations are linked to several diseases, including cancer. Lipids not only form lipid-lipid interactions but also interact with other biomolecules, including proteins, profoundly impacting each other.

Molecular dynamics (MD) simulations are a powerful tool to study the properties of cellular membranes and membrane-protein interactions on different timescales and at varying levels of resolution. Over the last few years, software and hardware for biomolecular simulations have been optimized to routinely run long simulations of large and complex biological systems. On the other hand, high-throughput techniques based on lipidomics provide accurate estimates of the composition of cellular membranes at the level of subcellular compartments. The community needs computational tools for lipidomics and simulation data effectively interacting to better understand how changes in lipid compositions impact membrane function and structure. Lipidomic data can be analyzed to design biologically relevant models of membranes for MD simulations. Similar applications easily result in a massive amount of simulation data where the bottleneck becomes the analysis of the data to understand how membrane properties and membrane-protein interactions are changing in the different conditions. In this context, we developed *LipidDyn*, an *in silico* pipeline to streamline the analyses of MD simulations of membranes of different compositions. Once the simulations are collected, *LipidDyn* provides average properties and time series for several membrane properties such as area per lipid, thickness, diffusion motions, the density of lipid bilayers, and lipid enrichment/depletion. The calculations exploit parallelization and the pipelines include graphical outputs in a publication-ready form. We applied *LipidDyn* to different case studies to illustrate its potential, including membranes from cellular compartments and transmembrane protein domains. *LipidDyn* is implemented in Python and relies on open-source libraries. *LipidDyn* is available free of charge under the GNU General Public License from https://github.com/ELELAB/LipidDyn.

## Introduction

Lipids are essential metabolites with crucial cellular functions and play a major role in most biological systems [1–3]. Lipid diversity, which depends on their chemical composition, is enormous and predicted to be in the range of hundreds of thousands [4,5], reflecting the variety of biological functions that lipids fulfil. Many different lipid species form the building blocks of cellular membranes [6]. Lipid compositions of cellular membranes can vary depending on the subcellular localization and are sensitive to cellular conditions and other factors [6]. Indeed, lipid alterations have been linked to different pathophysiological conditions, from cancer [5,7,8] to neurodegenerative diseases [9–11]. Lipid components of membranes are crucial determinants in the mechanism of action of several drugs [12]. Hence, targeting membrane lipids is becoming a possible therapeutic approach [13]. For instance, multidrug-resistant cancer cells present redistribution of phosphatidylserines from the inner leaflet of the plasma membrane, in which they mainly exist under physiological conditions, to the outer leaflet.

Lipids interact with other biomolecules, including proteins, and the two classes of biomolecules profoundly impact each other [14]. For example, lipids can influence protein dynamics and protein conformation [15]. On the other hand, membrane proteins can alter the biophysical properties of the lipids in the biological membranes [16].

Molecular dynamics (MD) simulations are a suitable tool to study the properties of cellular membranes and the membrane-protein interactions on different timescales and different levels of resolution, from coarse grain to all-atom representations [17]. The most commonly used physical models, i.e., force fields, for MD simulations, are Martini [18,19] and the ones in the CHARMM [20,21] and AMBER [22,23] families. These force fields cover most of the biologically relevant lipids and allow an accurate description of membranes including various lipid species and interaction with proteins. However, for complex systems, other force fields, as Slipids [24] or FUJI [25] may represent valuable alternatives. Recent developments in software and hardware for biomolecular simulations allow access to the microsecond-millisecond timescale of large and complex biological systems [26–28], such as lipid bilayers of heterogeneous composition [29].

A robust framework for MD simulations opens new venues to understand the complexity of biological membranes at the organelle level. One of the open challenges is how to design the lipid species and their ratio for the membrane models to use in simulations. On the experimental side, high throughput lipidomics provide elegant methodological solutions to profile lipids at the cellular [30–33] and organelle level [34,35]. In this context, we could envision using lipidomics data from assays performed in different cellular conditions on different subcellular fractions to design the bilayers to study with MD simulations. Similar applications will easily result in massive simulation data to analyze. Different tools calculate properties from MD simulations that can be compared to experimental observables from biophysical spectroscopies [36–42]. A bottleneck is making reproducible and simplifying the steps for analysis when several simulations should be analyzed in parallel. Pipeline engines can help in this regard.

In this context, we developed *LipidDyn*, an automated pipeline to streamline the analyses of MD simulations of membranes of different compositions. Our pipeline allows the estimate, in a non-time-consuming manner, of average properties and time series for different membranes. We also applied it to three cases studies as an example of *LipidDyn* applicability.

## Results

### Overview on LipidDyn

*LipidDyn* is a Python package for analyzing biophysical membrane properties and facilitating their interpretation in a non-time-consuming manner (**Figure 1**). *LipidDyn* allows to perform the analyses through an easy and practical Application Programming Interface (API) and implements full-fledged user programs accessible from the command line, including support for both analysis and plotting of the results. In this way, users can perform standard analyses of general interests on their molecular ensembles and write custom Python scripts that integrate several calculations seamlessly.

**Figure 1.**
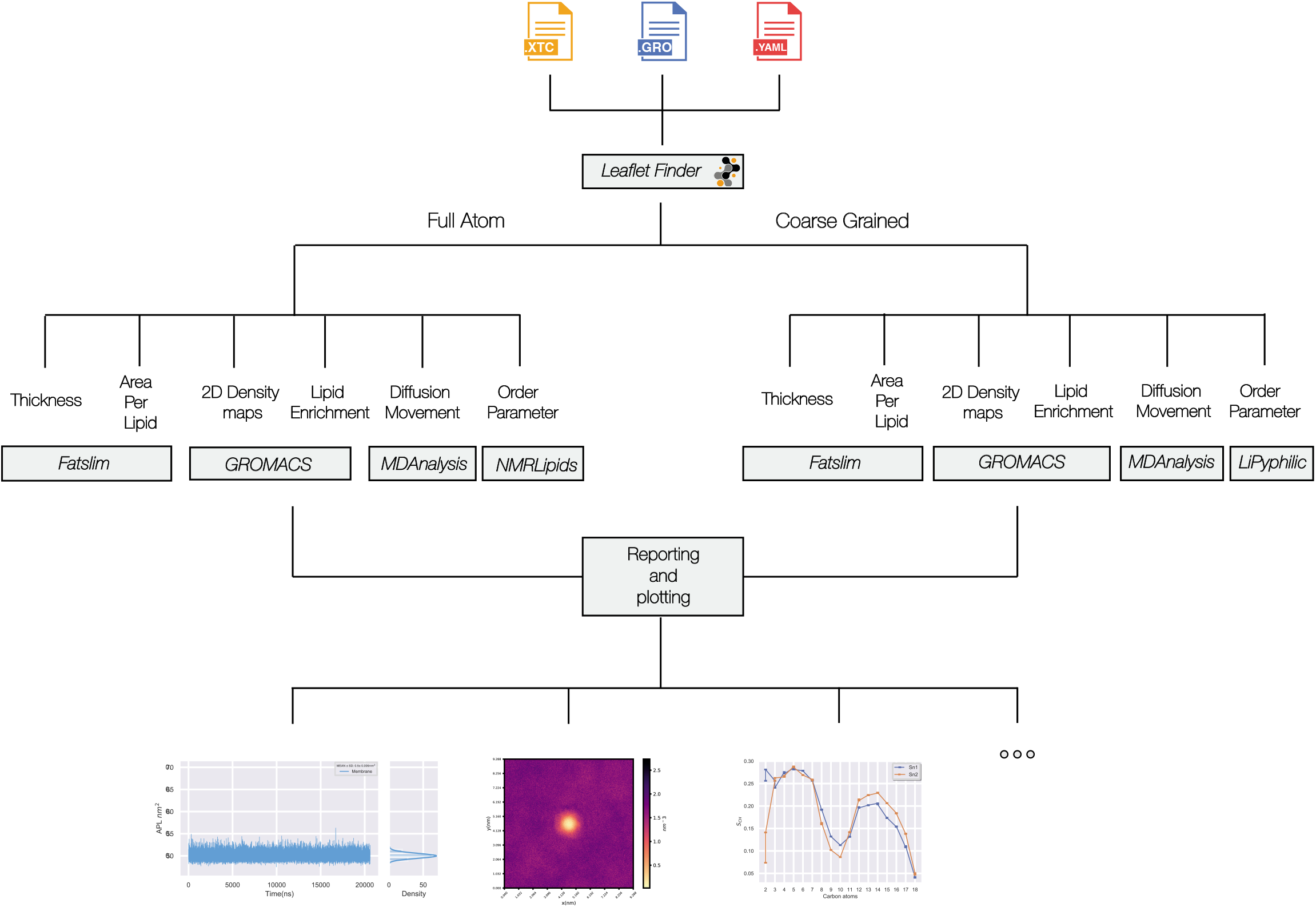
Overview of *LipidDyn*. The figure illustrates the workflow implemented in *LipidDyn* and its dependencies. The membrane is identified from the input files by the *MDAnalysis* tool *LeafletFinder*. Depending on the force field employed different methods are used for the analysis of choice.

*LipidDyn* is based on popular and well-maintained open-source packages, such as *MDAnalysis* [43], to handle trajectory files and other packages as back-ends [44].

While using the provided API ensures the highest degree of flexibility, it also requires extra programming work and expects the user to be familiar with Python packages. Nonetheless, the *LipidDyn* user scripts still allow for fine-tuning some aspects of the calculation either by command-line options or by using configuration files. This flexibility makes it possible to support simulations using different molecular mechanics physical models to represent the system under investigation. In addition, the workflow supports both the analyses of time-series and average properties. *LipidDyn* applies to both full-atom and coarse-grained topologies and trajectories. We include configuration files in the package for both short trajectories with all-atom (CHARMM36) and coarse-grained topologies (Martini).

We designed *LipidDyn* to process trajectory files in GROMACS format. It requires three input files: i) a configuration file (YAML format) including the definition (using the *MDanalysis* syntax) of each headgroup of the lipid species included in the system and the ratio of each lipid species to the total number of lipids in the system, ii) a topology file (.gro file), iii) a trajectory file (.xtc, .trr, or .gro).

*LipiDyn* handles trajectory files that contain many frames with data for large systems, including lipid bilayers and proteins. We tested *LipiDyn* with full-atom and coarse-grained trajectories of lipid bilayers and membrane proteins, including a different number of frames (10,000-200,000) and atoms/coarse-grained beads (10,000-50,000).

We focused on parameters that can be compared with experimental data.

*LipidDyn* includes different analyses, which can be performed independently or collectively. It consists of the calculation of i) membrane thickness, ii) area per lipid (APL), iii) two-dimensional (2D) lipid density maps, iv) lipid movements, v) lipid enrichment/depletion maps, vi) order parameter (**Figure 1**).

An essential prerequisite for analysis is the identification of the leaflets (defined as upper and lower leaflets) of the bilayer. In *LipidDyn*, we use the *LeafletFinder* class from *MDAnalysis* to identify which lipids belong to each leaflet, considering their headgroups or, in some cases, other atoms as representatives of each lipid molecule.

### Membrane thickness and area per lipid (APL)

The *FATSLiM* class performs APL and membrane thickness calculations [44]. The thickness is calculated for each lipid by using its neighborhood-averaged coordinates to remove the noise associated with fluctuations of lipid positions and then searching the neighbor lipids that belong to its opposite leaflet, using a cutoff distance (default: 6.0 nm). The thickness corresponds to the projection of the distance vector between each lipid and its neighbors in the opposite leaflet.

APL is estimated for each lipid by performing a neighbor search to identify its surrounding lipids in the leaflet and using them to compute a Voronoi tessellation. The implementation uses the Voronoi cell’s area to approximate the lipid’s APL. The program returns the upper and lower leaflet areas as the sum of the individual lipid areas and the membrane area as the average value of the two leaflet areas [44]. Compared to other existing tools [42,45–47], the computation with *FATSLiM* does not depend on the bilayer morphology, and it can accurately handle also vesicles. The user can visualize APL and thickness outputs with the *profiler* plotting tool included in the pipeline with options to customize the plot.

### Lipid density maps

The *Density* class performs lipid density calculation on both the upper and lower leaflet of the bilayer, providing 2D density maps. The class consists of a Python-based reimplementation of the density calculation algorithm provided by the *densmap* tool of GROMACS [48]. This algorithm divides the simulation box into a lattice of three-dimensional cells spanning a chosen dimension. Further, it calculates the time average of the number density of atoms across the plane of the remaining two dimensions. It visualizes local differences in lipid density with insights on lipid dynamics and system phase. The computed arrays are stored in *.dat* files for upper and lower leaflets. The user can visualize the outputs using the *dmaps* plotting tool to obtain 2D density maps.

### Lipid enrichment/depletion

The *Enrichment* class calculates the enrichment/depletion of each lipid species in specific regions of the bilayer, for example, around a membrane protein included in the system. For a given lipid species *L*, the class uses the *Density* class to compute the density map of the lipid *L* in the upper and lower leaflet averaged over the trajectory time. Then, the density map obtained is divided by the total number of lipids in the given leaflet. The resulting map is divided by the ratio of the lipid *L* in bulk.

The enrichment/depletion calculation is performed separately for the upper and lower leaflet of the bilayer. The user can visualize the outputs with the *dmaps* plotting tool to obtain 2D enrichment/depletion maps.

### Lipid movements maps

The *Movements* class provides graphical support for how each lipid moves along the X-Y plane of the bilayer. This analysis is useful to describe the motions of groups of lipids or even a single molecule of interest over trajectory time. The user can visualize the output with the *diffusion* tool to obtain 2D maps of lipid movements.

### Order Parameters

The *Order Parameter* class implements the calculation of the order parameter for the acyl chain tails of each lipid moiety. This analysis gives insight into the overall order of the lipid bilayer and the conformations that the acyl chains assume [49].

For the full-atom trajectories, the class includes a reimplementation of the algorithm in *NMRlipids* [https://github.com/NMRLipids] to calculate the carbon-hydrogen order parameter (*S_CH_*) of the acyl chains. For the coarse-grained trajectories, the class includes the Lipyphilic SCC() class [40] to calculate the carbon-carbon order parameter (*S_CC_*) of the acyl chains. For each lipid species, consecutive carbon atom pairs composing the *sn-1* and *sn-2* acyl chains are defined inside the configuration file. The *Order Parameter* class calculates *S_CH_* or *S_CC_* for the *sn-1* and *sn-2* acyl chains of each lipid species over trajectory time. The user can visualize the output with the *ordpar* plotting tool.

### Comparison with other tools

We compared the analyses provided by *LipidDyn* with other available tools (**Figure 2**) [40–42]. Each tool focuses on a group of analyses, with some classical ones in common, such as order parameter, thickness, and APL. *LiPyphilic* [40] and *Memsurfer* [42] provide mostly data on the bilayer’s geometrical properties as in the case of domain registration, z-positions, and z-angles calculations or membrane surface and curvature. On the other hand, *LOOS* [41] includes tools for analyzing membranes and accounts for an embedded protein. We noticed that most of these tools are either a suite of different scripts for analyses or a collection of Python classes to be imported. They do not provide complete workflows to streamline analysis collection and visualization and ensure reproducibility.

**Figure 2.**
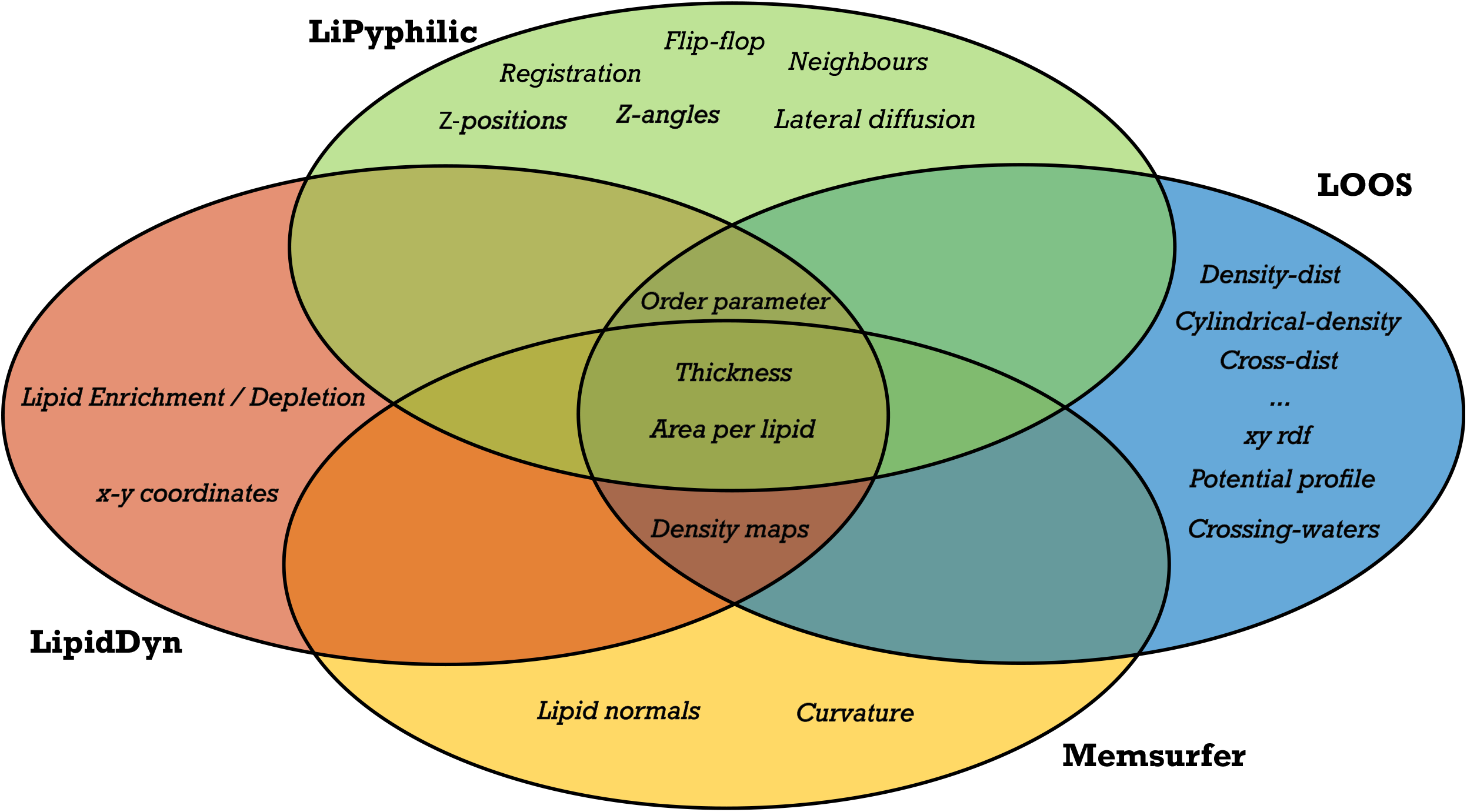
Comparison with other tools to analyze membrane simulations. The Venn diagram illustrates the comparison between the analyses covered by other available tools and *LipidDyn*. Most of the tools include the analysis of biophysical properties of lipid bilayers such as area per lipid, thickness, and order parameter. However, none of the tools currently cover all the possible analyses. Only *LipidDyn* has been designed as a workflow.

### Case study 1 - Analyses of the lipid behavior in ATG9A positive compartments

Autophagy is a catabolic process that mediates the degradation of cellular components by forming autophagosomes [50]. During autophagy, vesicles loaded with ATG9A translocate at the sites of the autophagosome formation, delivering lipids and proteins [51]. There is still scarce information about ATG9A-positive compartments, and structural studies on these compartments can bring new knowledge [52]. We performed liquid chromatography-mass spectrometry-based lipidomics of ATG9A-positive compartments immuno-isolated from amino acid–starved (i.e., autophagy-induced) HEK293A cells (see GitHub repository for tables summarizing the data). We then designed two models of membranes for full-atom MD simulations. The bilayers include a mixture of lipids designed from the composition of sphingomyelins quantified in the lipidomics data described above.

We used *LipidDyn* to analyze the MD simulations and compare them with a reference bilayer including only 1,2-dioleoyl-sn-glycero-3-phosphocholine (DOPC) (**Figure 3**). In particular, we calculated the time-series and average values for APL and thickness using the *FATSLiM* class. We also estimated the order parameter using the *Order Parameter* class of *LipidDyn* (**Figure 3**). APL and thickness are commonly used for the validation of bilayer MD ensembles. The average values of APL and thickness calculated from the DOPC trajectory are in good agreement with experimental values ~0.67 nm^2^ [53] and ~3.8 nm [54], respectively (**Figure 3A-B**). Our analysis shows that the presence of sphingomyelin is associated with a decrease of APL (average around 0.63 nm^2^) and a corresponding increase of thickness (average around 4.12 nm) (**Figure 3A-B**). The addition of cholesterol leads to higher lipid packing (average APL around 0.49 nm^2^), increased lipid chains order, and thicker bilayer (average thickness around 4.41 nm) (**Figure 3A-C**), showing a reorganization of the membrane structure. Our data are in agreement with experiments and simulations on the condensing effect of cholesterol on the membranes, which increases the order of the lipid packing and lowers the membrane permeability while maintaining membrane fluidity by forming liquid-ordered–phase lipid rafts with sphingolipids [55–58]. Our analyses shed light on the biophysical properties of the ATG9A-positive compartments upon autophagy induction, suggesting a certain degree of rigidity and packing dictated by the lipids that are enriched in these compartments.

**Figure 3.**
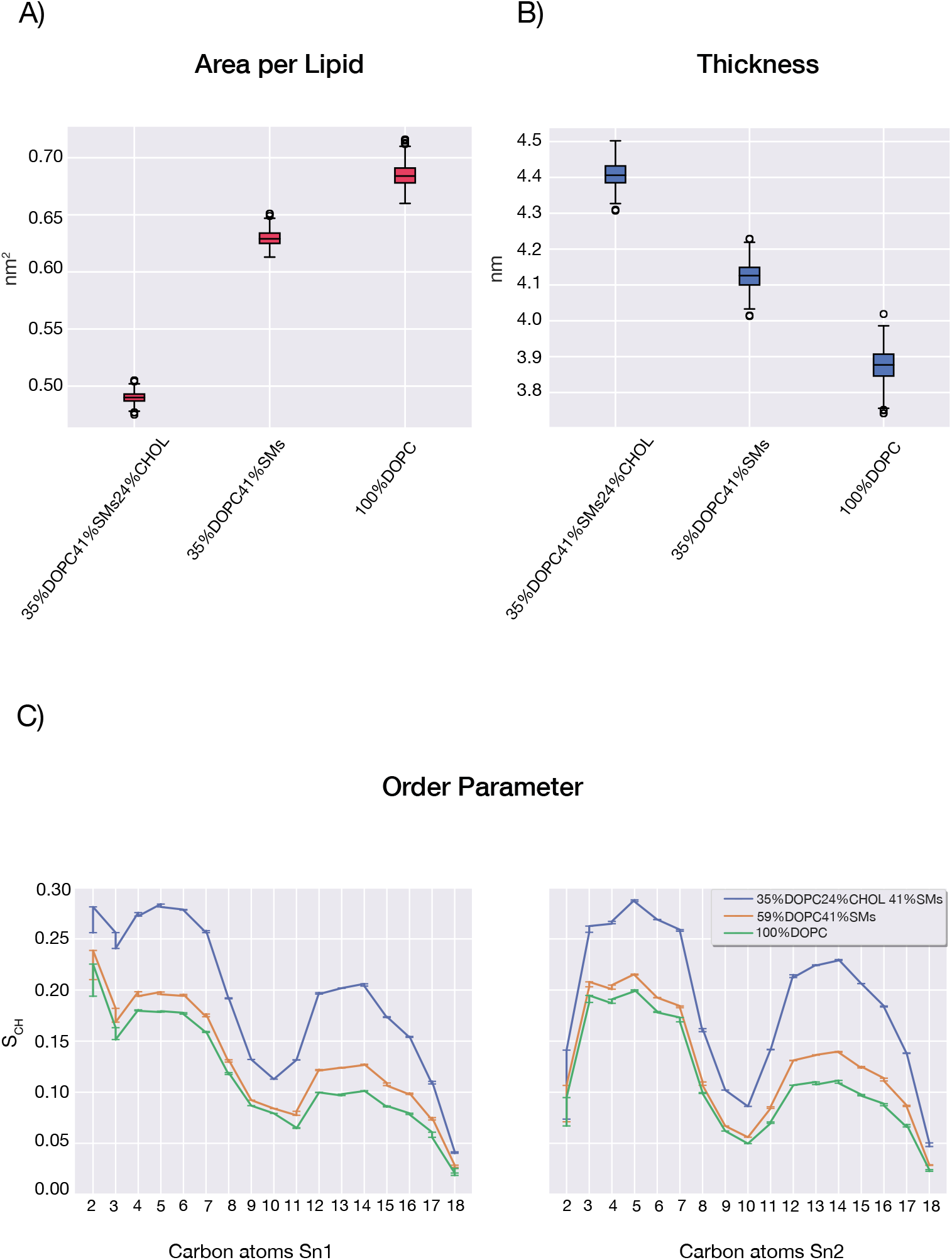
Analyses of MD simulations of ATG9A-positive compartments. A-B) Boxplot of the area per lipid and membrane thickness calculated for the full-atom simulations of the bilayers with lipid ratio DOPC 100%, DOPC 59% sphingomyelins (SMs) 41%, and DOPC 35% cholesterol (CHOL) 24% SMs 41%. C) Comparison of average order parameters for *sn1* and *sn2* acyl chain of DOPC between the presented systems. The ATG9A-positive compartments are associated with a decrease of area per lipid and an increase of thickness compared to the reference system. The addition of cholesterol leads to a higher lipid packing and lipid chains order and a thicker bilayer.

### Case study 2 - Lipidomics of single-organelle: structural properties of endoplasmic reticulum in HeLa cells

To analyze membranes from subcellular compartments in terms of structural and biophysical properties is essential not only for fundamental research but also for health-related applications [33,59]. Indeed, many diseases, including cancer, are associated with dysregulation of lipid metabolism [60].

We thus used immunoaffinity purification and mass spectrometry-based shotgun lipidomics to collect data from the endoplasmic reticulum (ER) of HeLa cells, quantifying 19 different lipid classes. We modeled a coarse-grained heterogeneous bilayer designed from the experimental lipidomics data, hereafter indicated as ER bilayer, and collected 10 μs MD simulation. The modeled ER bilayer includes 1,000 lipids for each leaflet. In detail, we had lipid species from the class of phosphatidylcholines (approximately 77%), a low concentration of cholesterol (approx. 6.3%), and sphingolipids (approx. 0.6%), in agreement with compositions previously reported [1]. As a comparison, we designed a coarse-grained bilayer with the same number of lipids per leaflet composed only by phosphatidylcholine (70% POPC) and higher cholesterol concentration (30%). With *LipidDyn*, we calculated the time-series and average values for APL and thickness using the *FATSLiM* class, and we estimated the average 2D lipid density using the *Density* class (**Figure 4**). The average values of the membrane thickness (average around 3.85 nm) and APL (average around 0.51 nm^2^) calculated from the reference POPC-cholesterol trajectory are in agreement with the known condensing effect of cholesterol [61], causing the thickening of the bilayer, reduction of APL, and ordering of the lipid tails (**Figure 4A-B**). Our analysis shows that the bilayer with a complex mixture and low content of cholesterol is associated with an increase of APL (average around 0.65 nm^2^) and a slight increase of thickness (average around 3.89 nm) (**Figure 4A-B**) in comparison to the POPC–cholesterol bilayer. The analysis of the average 2D lipid density maps shows regions of higher density in the POPC–cholesterol bilayer. In contrast, the ER bilayer shows a more uniform lipid density (**Figure 4C**), suggesting a more disordered lipid membrane. Our analyses shed light on the biophysical properties of ER bilayer, showing loose packing and low ordering of lipids that may reflect membrane dynamics involved in the functions of ER. Indeed, ER is at the beginning of the secretory pathway and at the level of its membrane happen the insertion and transport of newly synthesized proteins and lipids, as cholesterol which is synthesized at ER and then rapidly transported to other organelles [1,62,63].

**Figure 4.**
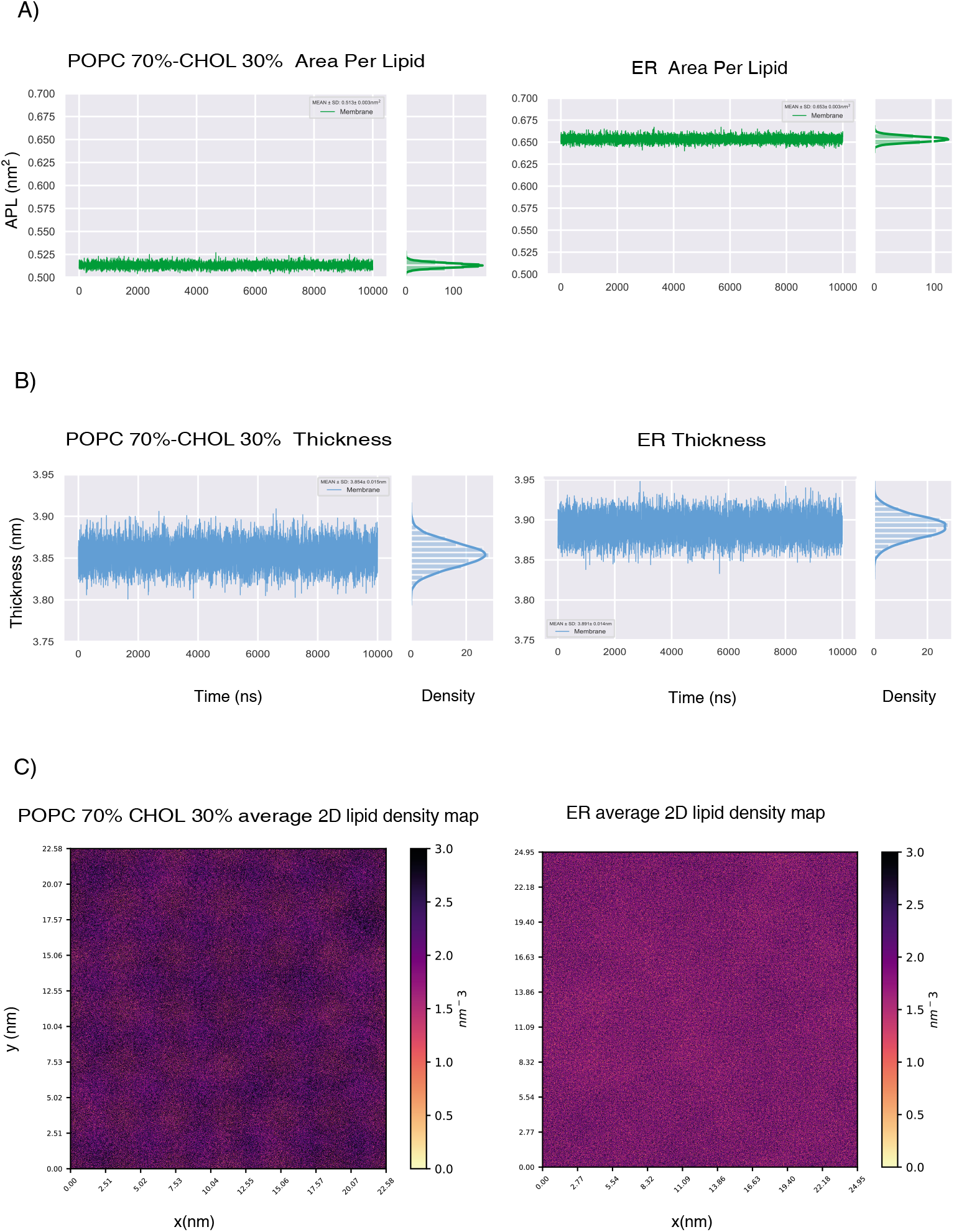
Analysis of coarse-grained MD simulations of the ER and POPC-cholesterol bilayers. A-B) Line plots of the area per lipid (panel A) and membrane thickness (panel B) calculated for the bilayer composed of phosphatidylcholine (POPC 70%) and cholesterol (CHOL 30%) and the bilayer designed from the lipidomics data of the endoplasmic reticulum (ER), which includes phosphatidylcholines (77.1%), CHOL (6.3%), sphingolipids (0.6%) and lipid species from other classes as phosphatidylethanolamine (6 %), phosphatidylinositol (5.8 %), ceramide (0.4 %), phosphatidylserine (0.3 %). Side distributions are also shown along with the line plots. C) Average 2D lipid density maps calculated for the upper leaflet of the bilayers. The ER bilayer is associated with an increase in the area per lipid and a more uniform lipid density than the POPC-cholesterol bilayer, suggesting loose packing and low ordering of lipids.

### Case study 3 - Study of the lipid interactions of the transmembrane emp24 domain 2 (p24) protein

The interactions of the transmembrane p24 protein with lipids [64–66] regulate its activity in the secretory pathway and vesicular trafficking [67]. The cytosolic part of the p24 transmembrane domain includes a sphingolipid-binding motif [64,68]. It has been shown that sphingolipids and ether lipids interact with the transmembrane helix of p24 and regulate p24’s cycling between ER and Golgi membranes, contributing to the early secretory pathway [65].

We used coarse-grained MD simulations to investigate if, with this approach, we can study the interaction of p24 with sphingomyelin, previously observed with full-atom MD simulations [64,65], and observe effects associated with the presence of cholesterol (**Figure 5**). We collected two 20 μs MD simulations of the transmembrane helix of p24, including residues 163-193, in the bilayer with lipid composition of phosphatidylcholine (70%-50% POPC), cholesterol (30%), and sphingomyelin (0%-20%). We used *LipidDyn* to investigate if p24 prefers interactions with specific lipid species in our systems. In particular, we used the *Density* class to calculate the 2D lateral density of the lipids and the *Enrichment* class to compute the enrichment-depletion map of each lipid species, considering the last μs of MD simulations (**Figure 5**). The analysis of the enrichment-depletion map in the POPC 70% cholesterol 30% system shows that cholesterol is enriched around the transmembrane helix of p24. This effect is more pronounced in the cytosolic leaflet of the membrane than the luminal one (**Figure 5A**). In sphingomyelin-rich membranes, cholesterol enrichment is reduced in the cytosolic leaflet, suggesting that sphingomyelin may affect cholesterol binding to p24 (**Figure 5B**). The map of sphingomyelin shows a diffused and more pronounced enrichment around the cytosolic portion of the transmembrane helix of p24, which is where the sphingolipid binding motif is located, than around the luminal part (**Figure 5C**). Although the limitations and approximations of the coarse-grained force field employed, especially in overestimating interactions [69], our analysis shed light on the interactions of the transmembrane helix of p24 with cholesterol and sphingomyelin, in agreement with previous data [64–66].

**Figure 5.**
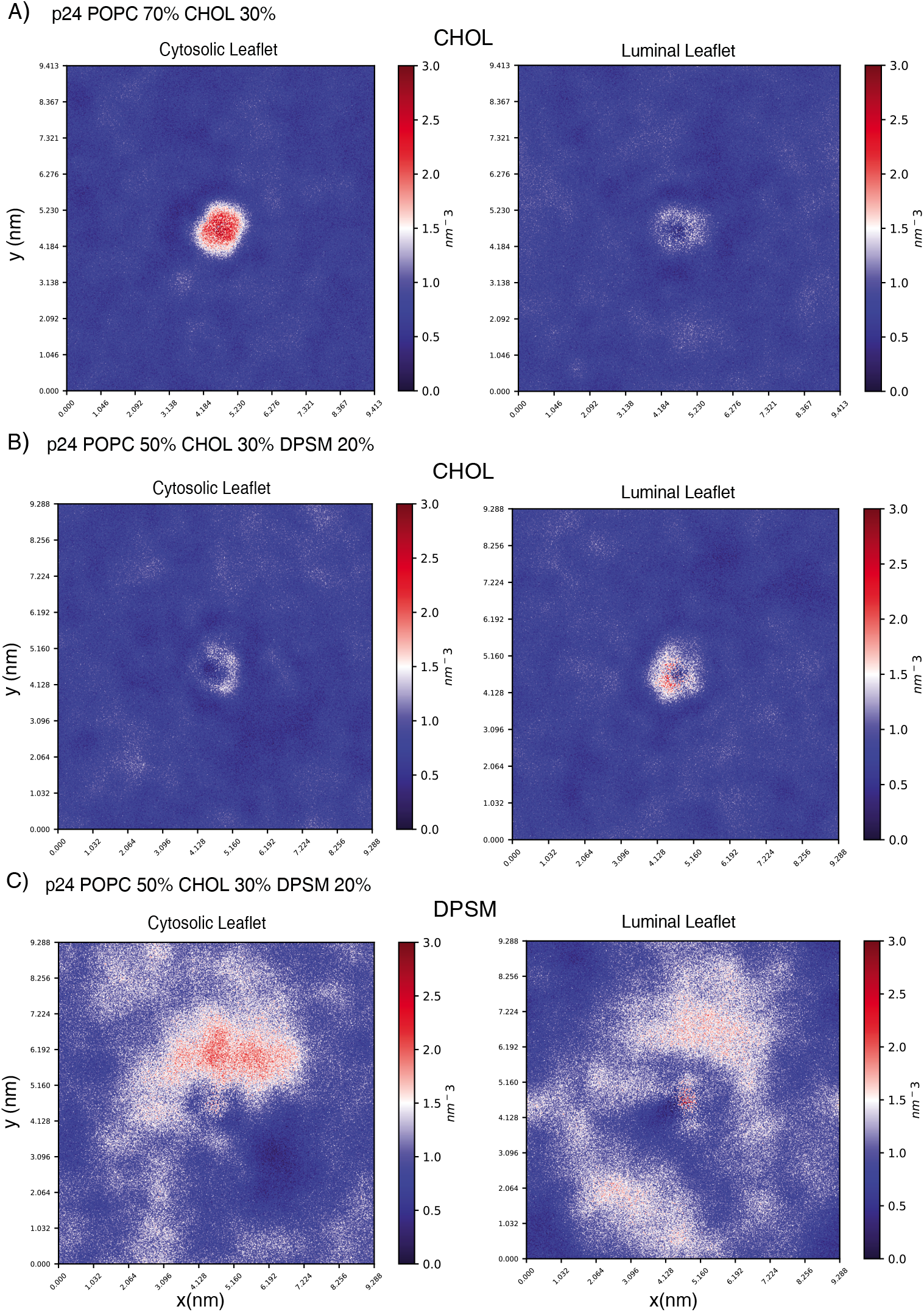
Analyses of coarse-grained MD simulations of the transmembrane domain of p24 embedded in different lipid bilayers. Enrichment-depletion map of A) CHOL in the cytosolic and luminal leaflet of the phosphatidylcholine (POPC 70%) and cholesterol (CHOL 30%) bilayer and of B) CHOL and C) sphingomyelin (DPSM) in the cytosolic and luminal leaflet of the phosphatidylcholine (POPC 50%), cholesterol (CHOL 30%) and sphingomyelin (DPSM 20%) bilayer. Our analysis shows a more pronounced sphingomyelin enrichment around the cytosolic part of the transmembrane domain of p24, which includes the sphingolipid binding motif. The inclusion of sphingomyelin in the bilayer affects the binding of cholesterol to p24 in the cytosolic leaflet.

## Materials and Methods

### Lipidomics of the endoplasmic reticulum of HeLa cells

The post-nuclear lysate of HeLa cervical cancer cells was prepared as previously described [70]. 300 μl post-nuclear lysate was incubated with 0.6 μg/ml rabbit anti-calnexin antibody (ab22595, Abcam) for 45 minutes and for an additional one hour after addition of magnetic microbeads conjugated to anti-rabbit IgG (25 μl, 30-048-602; Miltenyi Biotec) to purify the endoplasmic reticulum. The endoplasmic reticulum was then captured on an MS column (130-042-201; Miltenyi Biotec) mounted on an OctoMacs magnetic separator (130-042-108; Miltenyi Biotec) and eluted after washing and demounting of the column. The eluted endoplasmic reticulum was pelleted by centrifugation for 20 minutes at 21,100 g and resuspended in 200 μl 155 mM ammonium bicarbonate. The entire purification procedure was performed at 4 °C. Lipid extraction and mass spectrometry-based lipidomics analysis was carried out as previously described [71].

### Lipidomics of ATG9A-positive lipid compartments

HEK293A cells were cultivated in a full medium composed of DMEM supplemented with 10% FCS and 4 mM l-glutamine as described in [51]. We performed the immunoisolation of ATG9A-positive compartments from HEK293 cells as previously described [51]. We carried out metabolite extraction by fractionating the cell samples into pools of species with similar physicochemical properties, using combinations of organic solvents. Cell pellets were resuspended in cold water and briefly mixed. Proteins were precipitated from the lysed cells by adding methanol. The samples were spiked with chloroform after vortex mixing. The extraction solvents were spiked with metabolites not detected in unspiked cell extracts used as internal standards. We incubated the samples at −20 °C for 30 minutes and collected two different phases after a vortex step. Cell extracts were mixed with water (pH 9), and after brief vortexing, the samples were incubated for 1 hour at −20 °C. After centrifugation at 16,000 x g for 15 minutes, the organic phase was collected. We reconstituted the dried extracts in acetonitrile/isopropanol (50:50), resuspended them for 10 minutes, centrifuged (16,000 x g for 5 minutes), and transferred them to vials for UPLC-MS analysis.

We used two quality controls described in [72]. Randomized duplicate sample injections were performed. We performed pre-processing, normalization, and statistical analysis with TargetLynx application manager for MassLynx 4.1 software (Waters Corp., Milford, USA). The processing was executed using a set of predefined retention time, mass-to-charge ratio pairs, *Rt-m/z*, corresponding to metabolites included in the analysis. The ion chromatograms were denoised and peak-detected with a mass tolerance window of 0.05 Da. For each sample injection, a list of chromatographic peak areas was generated. We used representative MS detection curves to identify the metabolites using internal standards. The normalization factors were calculated for each metabolite by dividing their intensities in each sample by the recorded intensity of an appropriate internal standard in that same sample, as described in [72]. Statistical analysis included principal component analysis, Shapiro-Wilk test, Student t-test, and Wilcoxon-signed-rank test.

### From lipidomics to MD design

We designed a Python script (see GitHub) to analyze the ATG9A-positive lipidomic dataset. We quantified the average lipid concentration from the raw data among the two starved samples. To design the lipid composition for the MD simulation, we associated the sphingomyelin species identified in the lipidomic dataset with those available in the CHARMM36 force field, querying the *LipidDyn* internal database.

We designed a Python script (see GitHub) to analyze the lipidomics dataset of the endoplasmic reticulum from HeLa cells at the level of classes of lipids and to design the lipid composition of the bilayer for the coarse-grained MD simulations using Martini force field. In addition, we have been curating a more general dictionary to link lipid species to the corresponding available molecules in all-atom and coarse-grained force fields to assist the design of the lipid composition for MD simulations.

### MD simulations

We used CHARMM-GUI Membrane Builder [73,74] to build the systems for simulations. The MD simulations were carried out using GROMACS [48].

We used CHARMM36 force field [21] and the TIP3P water model adjusted for CHARMM force fields[75] for all-atom MD simulations and Martini 2 for coarse-grained simulations [76]. Each bilayer system was built in a rectangular box in the x and y dimension. The water thickness (minimum water height on top and bottom of the system) was set between 25 Å and 35 Å to ensure that the two layers of water molecules were sufficient to avoid artificial contacts between the image boxes. Each system was simulated for different timescales ranging from 0.5 to 20 μs. Periodic boundary conditions were applied in all three dimensions. More details, including the preparation steps, are reported in the readme files in the GitHub or OSF repositories for each set of simulations.

## Availability and future perspective

The package and test cases are available at https://github.com/ELELAB/LipidDyn.

*LipidDyn* has been released in its first version to provide a well-organized workflow for analyses of lipid and protein-lipid simulations and streamline cases where many bilayers with different compositions need to be analyzed in parallel. We selected the most important parameters that often need to be scrutinized, with emphasis on supporting properties that can also be experimentally determined. Nevertheless, *LipidDyn* at the moment does not cover all the available portfolio of analyses of structural and biophysical properties that can be applied to membrane simulations. In the future, we will widen the range of analyses supported by the package and also find new visualization solutions. For example, we will include tools to calculate the shapes and curvatures of lipid membranes. In terms of protein-lipid interaction, we will implement analyses of occurrence, maximal occupancy and time life of contacts along the simulation time. We will provide *LipidDyn* outputs compatible with Pyinteraph2 [77,78] so that protein-membrane simulations can be analyzed using methods from graph theory.

Furthermore, we plan to include in *LipidDyn* automated support to convert the information in the processed lipidomics data to design models of membranes for MD simulations that resemble experimentally observed lipid compositions. Our focus is to provide a tool in *LipidDyn* that includes a dictionary to automatize the mapping and conversion of the lipid species in the lipidomics datasets to the corresponding molecules for which parameters are available in the commonly used force fields for MD simulations.

## Acknowledgments

E.P. group is supported by Danmarks Frie Forskningsfond, Natural Science, Project 1 (102517), NovoNordisk Fonden Bioscience and Basic Biomedicine (NNF20OC0065262) and Andreas og Grethe Gullev Hansens Fond (to M.L.). M.J. is supported by NovoNordisk Distinguished Investigator Grant - Endocrinology and Metabolism (054296). M.J. and E.P. groups are part of the Center of Excellence for Autophagy, Recycling, and Disease (CARD), which is supported by Danmarks Grundforskningsfond (DNRF125). K.M. is supported by Danmarks Frie Forskningsfond, Sapere Aude (6108–00542B). D.J. and S.A.T. were supported by The Francis Crick Institute which receives its core funding from Cancer Research UK (FC001187, FC001999), the UK Medical ResearchCouncil (FC001187, FC001999). This research was funded in whole, or in part, by the Wellcome Trust (FC001187, FC001999).

The calculations described in this paper were performed thanks to DECI-PRACE 17th to ML for calculations on Archer2 (UK). Part of the calculations has been carried out with resources available at the DeiC National Life Science Supercomputer (Computerome2) at the Technical University of Denmark. The authors would like to thank Valeria Zoni and Stefano Vanni for sharing the protocol and data to test our implementation of the *Enrichment* class on previously published trajectories. Moreover, the authors would like to thank Matteo Arnaudi, Ludovica Beltrame, Matteo Orlandi, and Mattia Utichi for testing installation and case studies on different architectures. We also thank Dr. Mesut Bilgin and the Lipidomics Core Facility of DCRC for making instruments and materials available for lipidomics experiments.

